# Validation of a SARS-CoV-2 spike protein ELISA for use in contact investigations and sero-surveillance

**DOI:** 10.1101/2020.04.24.057323

**Authors:** Brandi Freeman, Sandra Lester, Lisa Mills, Mohammad Ata Ur Rasheed, Stefany Moye, Olubukola Abiona, Geoffrey B. Hutchinson, Maria Morales-Betoulle, Inna Krapinunaya, Ardith Gibbons, Cheng-Feng Chiang, Deborah Cannon, John Klena, Jeffrey A. Johnson, Sherry Michele Owen, Barney S. Graham, Kizzmekia S. Corbett, Natalie J. Thornburg

## Abstract

Since emergence of SARS-CoV-2 in late 2019, there has been a critical need to understand prevalence, transmission patterns, to calculate the burden of disease and case fatality rates. Molecular diagnostics, the gold standard for identifying viremic cases, are not ideal for determining true case counts and rates of asymptomatic infection. Serological detection of SARS-CoV-2 specific antibodies can contribute to filling these knowledge gaps. In this study, we describe optimization and validation of a SARS-CoV-2-specific-enzyme linked immunosorbent assay (ELISA) using the prefusion-stabilized form of the spike protein [1]. We performed receiver operator characteristic (ROC) analyses to define the specificities and sensitivities of the optimized assay and examined cross reactivity with immune sera from persons confirmed to have had infections with other coronaviruses. These assays will be used to perform contact investigations and to conduct large-scale, cross sectional surveillance to define disease burden in the population.

## Background

Since the emergence of SARS-CoV-2 in late 2019, there has been a need to identify transmission patterns, determine prevalence, and determine the true burden of disease and case fatality rates. Case fatality rates have differed between countries [2–4]. Defining these rates has been difficult due to various factors including f asymptomatic infection [5–7] and limits to molecular testing capacity. Several reports have confirmed that most patients with established SARS-CoV-2 infections mount serum antibody responses specific to viral proteins [8–10]. Because seroconversion may not occur for 1-3 weeks after symptom onset, antibody testing may have limited utility for diagnosis of acute infection. However, detection of anti-SARS-CoV-2 serum antibody responses can be used to define transmission chains in contact investigations. Additionally, in cross -sectional serosurveillance studies, antibody assays can be used to define the burden of disease and be used for more accurate calculations of case fatality rates.

## Objectives

In this manuscript we describe assay optimization and validation of a SARS-CoV-2 spike protein ELISA. We intend to use this assay in contact investigations to identify individuals who had a SARS-CoV-2 infection without prior molecular diagnostic confirmation. We plan to use these assays to study the natural history of infection to determine the percent of individuals with a range of disease severity who mount serum antibody responses against the virus. Finally, we plan to use these assays in large-scale serosurveillance to better define, on a population basis, the number of individuals who may have had COVID-19, including those with mild and asymptomatic infections. These studies will be essential for characterizing transmission, identifying the true burden of disease, existence of population-wide serum antibodies, and calculating accurate case fatality rates. To this end, we needed to define the parameters that maximized the sensitivity and specificity of anti-SARS-CoV-2 antibody-detection assays.

## Study design

### Sera collection

True negative sera were collected between 2011 and 2019 from 519 adults who were healthy (2016-2019, n = 377) or collected from suspected hanta virus patients (2016-2019, n = 101), HIV (2011-2012, n = 21), hepatitis B virus (2011-2012, n = 10), or HCV positive (2011-2012, n = 10). Sera from hantavirus sera were used because cases had respiratory virus infections and found to be hantavirus negative. They, therefore, represent negative controls with recent respiratory virus infections. Convalescent sera from PCR+ COVID-19 cases were collected at day 10 post-symptom onset or later (n = 99). Additionally, acute and convalescent paired sera from PCR-confirmed commonly circulating coronavirus (229E, NL63, OC43, and HKU1)- infected patients were collected as previously described [11].

### Ethics

All serum specimens were de-identified. The investigation was determined to constitute non-human subjects research by CDC National Center for Immunization and Respiratory Diseases (project 0900f3eb81b07602).

### ELISA

The pre-fusion stabilized ectodomain of SARS-CoV-2 spike (S) was expressed in suspension adapted HEK-293 cells as previously described [1]. Coating conditions were optimized by antigen dilution and testing with convalescent sera collected from two COVID-19 patients at days 15 and 23 post symptom onset. Coating concentrations ranging from 0.019– 5 μg / ml were tested. Antigen was diluted in PBS to 0.15 μg / ml. The top half of Immulon 2 HB (Thermo Fischer) 96 well-plates were coated with antigen, and the bottom half with PBS overnight at 2-8°C in a humidified chamber. The next day, 5 X Stabilcoat blocker (Surmodics) was diluted 1:1 in PBS. Plates were washed 3 times with PBS tween 20 (PBS-T)diluted from a 10 X stock pH 7.4 – 7.6 (KD Medical catalog # 0125) using a Biotek plate 405 washer and were blocked with 2.5 2.5 X Stabilcoat blocking buffer at 37° C for one hour. During the hour blocking plates, sera were diluted 1:25 in serum diluent (PBS-T / 5% skim milk), including positive and negative controls. After blocking, plates were washed 3 Xs with PBS-T, and 100 μl serum diluent was added to all wells. Thirty three point three μl serum diluted to 1:25 was added to rows A and E and mixed by pipetting. Four-fold serially dilutions were achieved by pipetting 33.3 ul diluted sera down the plates in rows A-D, and E-H, discarding 33.3 ul from rows D and H after dilutions. Plates were incubated at 37°C in a humidified chamber for one hour and washed 3 Xs with PBS-T. HRP-conjugated goat anti-human antibodies (KPL) diluted at 1:2000 in serum diluent were added to washed plates and incubated at 37°C in a humidified chamber for 1 hour. During secondary antibody incubation, ABTS peroxidase substrate system (KPL) components were brought to room temperature and prepared according to the manufacturer’s instructions. Plates were washed 3 Xs with PBS-T, 100 μl prepared substrate was added to each well, and were incubated for 30 minutes at 37°C. During substrate incubation, ABTS stop solution was prepared according to the manufacturer instruction. After 30 minutes, 100 μl prepared stop solution was added to each well, and each plate was read at 405 and 490 nm using a PerkinElmer Victor XV plate reader. Final background corrected ODs were calculated as 490-405, antigen coated – PBS coated well for each dilution of each specimen. Statistics were performed using GraphPad Prism software (7.04).

## Results

Coating with 0.15 μg / ml of antigen was found to produce a saturated signal in convalescent sera that titrated with serum dilution (data not shown). Receiver operator characteristic (ROC) analyses of spike protein ELISAS were conducted using anti-human pan Ig, anti-human IgG, and anti-human IgM secondary antibodies (Figure 1A). Two dilutions, 1:100 (Figure 1A) and 1:400 (Figure 1B), were analyzed using the pan-Ig secondary antibody. At a background corrected OD of 0.4 with serum diluted 1:100, specificity was 99.3% (confidence interval 98.32 – 99.88%) and sensitivity was 96% (confidence interval 89.98 – 98.89%) (Figure 1A). Using anti-IgG secondary antibodies at the same OD cutoff, sensitivity was 94.74% (confidence interval 85.38 – 98.9%), and 76.19% sensitivity (confidence interval 63.79 – 86.02%) using anti-IgM secondary antibody (Figure 1A). Diluting sera to 1:400 reduced the sensitivity to 86.7% (Figure 1B). These data indicate diluting sera at 1:100 and using pan Ig secondary antibody maximizes specificity and sensitivity.

**Figure 1.**
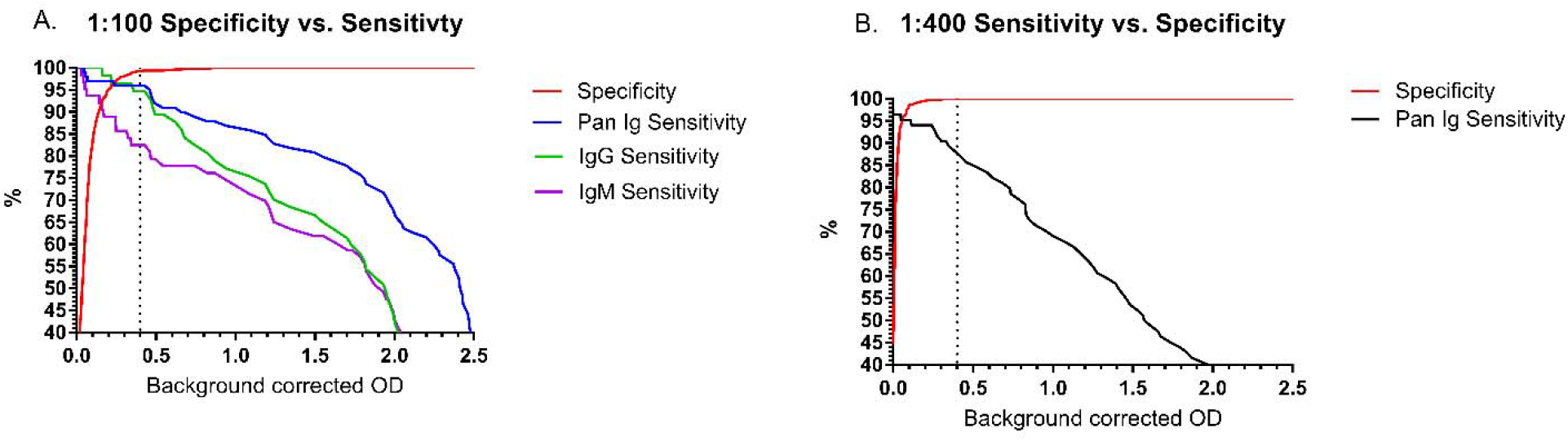
ROC analysis of 519 true negative and 99 true positive sera. A. Specificity was calculated at a 1:100 dilution using pan-Ig secondary (red). Sensitivities were calculated with sera diluted to 1:100 using anti-Pan Ig (blue), anti-IgG (green) or anti-IgM (purple) secondary antibodies. B. Results were analyzed with sera diluted to 1:400 using anti-Pan Ig secondary antibody.

To examine potential cross reactivity with common coronavirus sera, healthy donor sera, and convalescent immune sera SARS-1 (n =4), MERS-CoV (n = 1), SARS-CoV-2 (n = 9), NL63 (n = 8), OC43 (n = 17), and HKU1 (n = 14), and 229E (n = 3) patients were tested using the same ELISA conditions and pan-Ig secondary antibody. SARS-1 and MERS-CoV sera exhibited cross reactivity to SARS-CoV-2 spike protein, while NL63, OC43, HKU1, and 229E sera did not (Figure 2A). To examine rising cross reactivity that may be present in sera from confirmed common coronavirus cases, acute and convalescent paired sera were examined. There were not rising signals between acute and convalescent timepoints in any of 8 paired NL63 or 2 pairs of 229E sera (Figure 2B and E). There were 2 of 17 OC43 paired specimens that exhibited rising signals between acute and convalescent timepoints, though still below the set cutoff (Figure 2C). Similarly, there were 2 of 14 HKU1 paired specimens that exhibited rising signals between acute and convalescent timepoints, though still below the set cutoff (Figure 2D). These data indicate that there may be significant cross reactivity to MERS-CoV and SARS-1 sera, though prior infections with those viruses are extremely rare and should not contribute to background reactivity in large scale serosurveillance or in contact investigations. More common coronavirus infections do exhibit some cross reactivities, however they are below the limits of detection.

**Figure 2.**
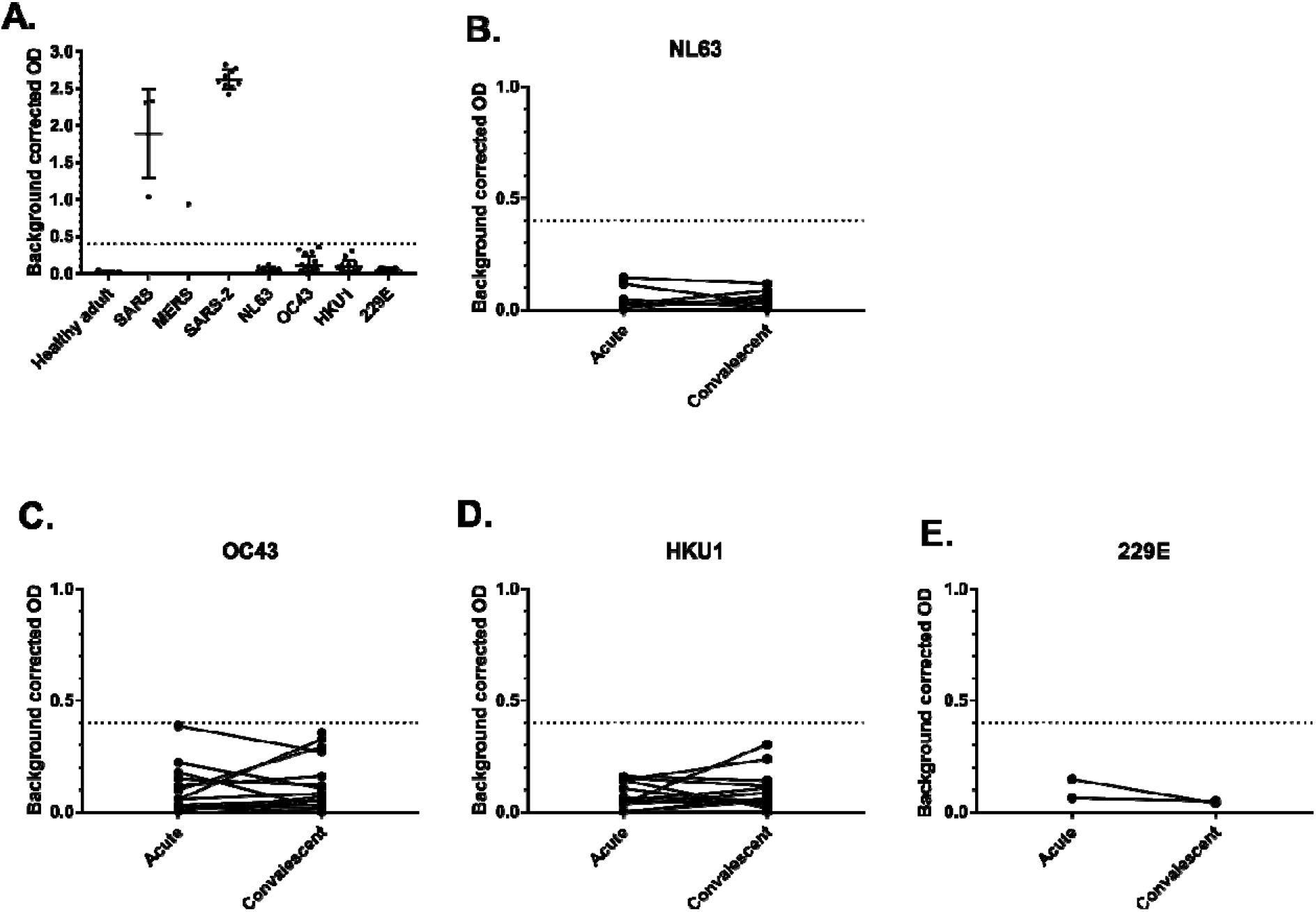
Examination of coronavirus sera cross reactivity to SARS-CoV-2 ELISA at a dilution of 1:100. The dotted line represents the OD cutoff established by ROC analysis. A. Healthy adult, SARS1 n =4, MERS n = 1, SARS-2 n = 9, NL63 n = 8, OC43 n = 17, HKU1 n = 14, and 229E (n = 3). B – E Examination of rising signals between acute and convalescent from PCR cases of B.NL63 (n = 8),C. OC43 (n = 17), D. HKU1 (n = 14), and E. 229E (n = 2).

## Conclusions

We have optimized an ELISA against SARS-CoV-2 spike protein that is sensitive and specific. Diluting sera 1:100 and use of goat anti-human Pan Ig secondary antibody maximized sensitivity and specificity of the assay. While we did see some cross reactivity with SARS1 and MERS-CoV sera, there was minimal cross reactivity using sera against commonly circulating coronaviruses. With specificity >99%, and a sensitivity of 96%, this assay can be used to identify prior SARS-CoV-2 infections without molecular diagnostic confirmation. This assay can be adjusted for use in higher throughput serosurveys and in combination with confirmatory testing could be used in an algorithm for informing individuals of prior infection.

## Acknowledgements

We thank the Bioexpression and Fermentation Facility, University of Georgia for large scale production of the protein. The work was completed by Dr. Neda Maleki, Dr. Chris Keggi, Dr. L. Michelle Lewis. We also thank Dr. Brian Harcourt for inactivation support, Jarad Schiffer for assay design input, and the CDC Divisions of Viral Hepatitis and HIV/AIDS Prevention for providing samples used in cross-reactivity testing. This work was supported in part by intramural funding from the National Institute of Allergy and Infectious Diseases.

## Disclaimers

The findings and conclusions in this report are those of the author(s) and do not necessarily represent the official position of the Centers for Disease Control and Prevention. Names of specific vendors, manufacturers, or products are included for public health and informational purposes; inclusion does not imply endorsement of the vendors, manufacturers, or products by the Centers for Disease Control and Prevention or the US Department of Health and Human Services.

